# *ESR1*, *WT1*, *WNT4, ATM* and *TERT* loci are major contributors to uterine leiomyoma predisposition

**DOI:** 10.1101/291237

**Authors:** Niko Välimäki, Heli Kuisma, Annukka Pasanen, Oskari Heikinheimo, Jari Sjöberg, Ralf Bützow, Nanna Sarvilinna, Hanna-Riikka Heinonen, Jaana Tolvanen, Simona Bramante, Tomas Tanskanen, Juha Auvinen, Terhi Piltonen, Amjad Alkodsi, Rainer Lehtonen, Eevi Kaasinen, Kimmo Palin, Lauri A. Aaltonen

**Author notes:** These authors contributed equally to this work (Niko Välimäki, Heli Kuisma). Correspondence to: Lauri A. Aaltonen, Department of Medical and Clinical Genetics, Biomedicum Helsinki, P.O. Box 63 (Haartmaninkatu 8), FI-00014 University of Helsinki, Finland.

## Abstract

Uterine leiomyomas (ULs) are benign tumors that are a major burden to women’s health. A genome-wide association study on 5,417 UL cases and 331,791 controls was performed, followed by replication of the genomic risk in two cohorts. Effects of the identified risk alleles were evaluated in view of molecular and clinical features.

Five loci displayed a genome-wide significant association; the previously reported *TNRC6B,* and four novel loci *ESR1 (ERα)*, *WT1*, *WNT4,* and *ATM*. The sixth hit *TERT* is also a conceivable target. The combined polygenic risk contributed by these loci was associated with *MED12* mutation-positive tumors. The findings link genes for uterine development and genetic stability to leiomyomagenesis. While the fundamental role of sex hormones in UL aetiology has been clear, this work reveals a connection to estrogen receptor alpha on genetic level and suggests that determinants of UL growth associated with estrogen exposure have an inherited component.

## INTRODUCTION

Uterine leiomyomas (ULs), also known as fibroids or myomas, are benign smooth muscle tumors of the uterine wall. They are extremely common; approximately 70% of women develop ULs before menopause^1^. The symptoms, occurring in one fifth of women, include excessive menstrual bleeding, abdominal pain and pregnancy complications^1^. In most cases, durable treatment options are invasive^2^. ULs cause a substantial human and economic burden, and the annual cost of treating these tumors has been approximated to be as high as $34 billion in the United States, higher than the combined cost of treating breast and colon cancer^3^.

Earlier studies have indicated strong genetic influence in UL susceptibility based on linkage^4^, population disparity^5^ and twin studies^6^. The most striking UL predisposing condition thus far characterized is hereditary leiomyomatosis and renal cell cancer (HLRCC) syndrome, caused by high-penetrance germline mutations in the *Fumarate hydratase* (*FH*) gene^7,8^. Genome-wide association studies (GWAS) have proposed several low-penetrance risk loci but few unambiguous culprit genes have emerged. Cha *et al.* reported loci in chromosome regions 10q24.33, 11p15.5 and 22q13.1 based on a Japanese patient cohort^9^. The 11p15.5 locus - near the *Bet1 golgi vesicular membrane trafficking protein like* (*BET1L*) gene - was later replicated in European Americans^10^. The 22q13.1 locus has been replicated in Caucasian, American and Saudi Arabian populations suggesting *trinucleotide repeat containing 6B* (*TNRC6B*) as a possible target gene^10–12^. Further UL predisposition loci have been suggested at 1q42.2 and 2q32.2 by Zhang *et al.*^13^ and, at 3p21.31, 10p11.21 and 17q25.3 by Eggert *et al.*^14^ A recent work reported *cytohesin 4* (*CYTH*4) at 22q13.1 as a novel candidate locus in African Americans^15^. While multiple loci and genes have been implicated through these valuable studies it is not straightforward to connect any of them mechanistically to UL development.

Most ULs show somatic site-specific mutations at exons 1 and 2 of the *mediator complex subunit 12* (*MED12*) gene^16,17^. These observations together with further scrutiny of driver mutations, chromosomal aberrations, gene expression, and clinicopathological characteristics have lead to identification of at least three mutually exclusive UL subtypes; *MED12* mutant, *Fumarate Hydratase* deficient, as well as *HMGA2* overexpressing lesions^18^.

Here we report the most powerful GWAS on uterine leiomyoma to date, and novel genome-wide significant UL susceptibility loci with plausible adjacent culprit genes including *Estrogen Receptor 1* (*ESR1 or ERα*), *Wilms Tumor 1* (*WT1*), *Wnt Family Member 4* (*WNT4*) and *ATM Serine/Threonine Kinase* (*ATM*). Genome-wide significant associations were observed also at the previously reported locus in *TNRC6B,* and after meta-analysis at a gene poor locus 13q14.11. *Telomerase Reverse Transcriptase* (*TERT*) locus narrowly missed genome-wide significance, but *TERT* appeared as a highly plausible UL predisposition gene supported by overwhelming evidence in many other tumor types^19^. We compiled the weight of these susceptibility loci into a polygenic risk score and replicated the UL association in an independent cohort of Finnish origin. Finally, we investigated the risk alleles’ association to clinical features, molecular UL subtypes, telomere length, gene expression and DNA methylation.

## METHODS

### Genome-wide association study

Fig. 1 provides an outline of the four stages that were implemented. The discovery stage GWAS was based on UK Biobank (UKBB) genotypes: a Global Biobank Engine (GBE) query was used to test association between 5,417 UL cases and 331,791 controls of Caucasian ancestry. The second stage meta-analysis utilized the genome-wide summary statistics from UKBB and the Helsinki cohort of 457 UL cases and 15,943 controls. The observations were replicated using 459 UL cases and 4,943 controls from the Northern Finland Birth Cohort (NFBC). More details on the GWAS materials and methods are given in Supplementary Methods.

**Figure 1:**
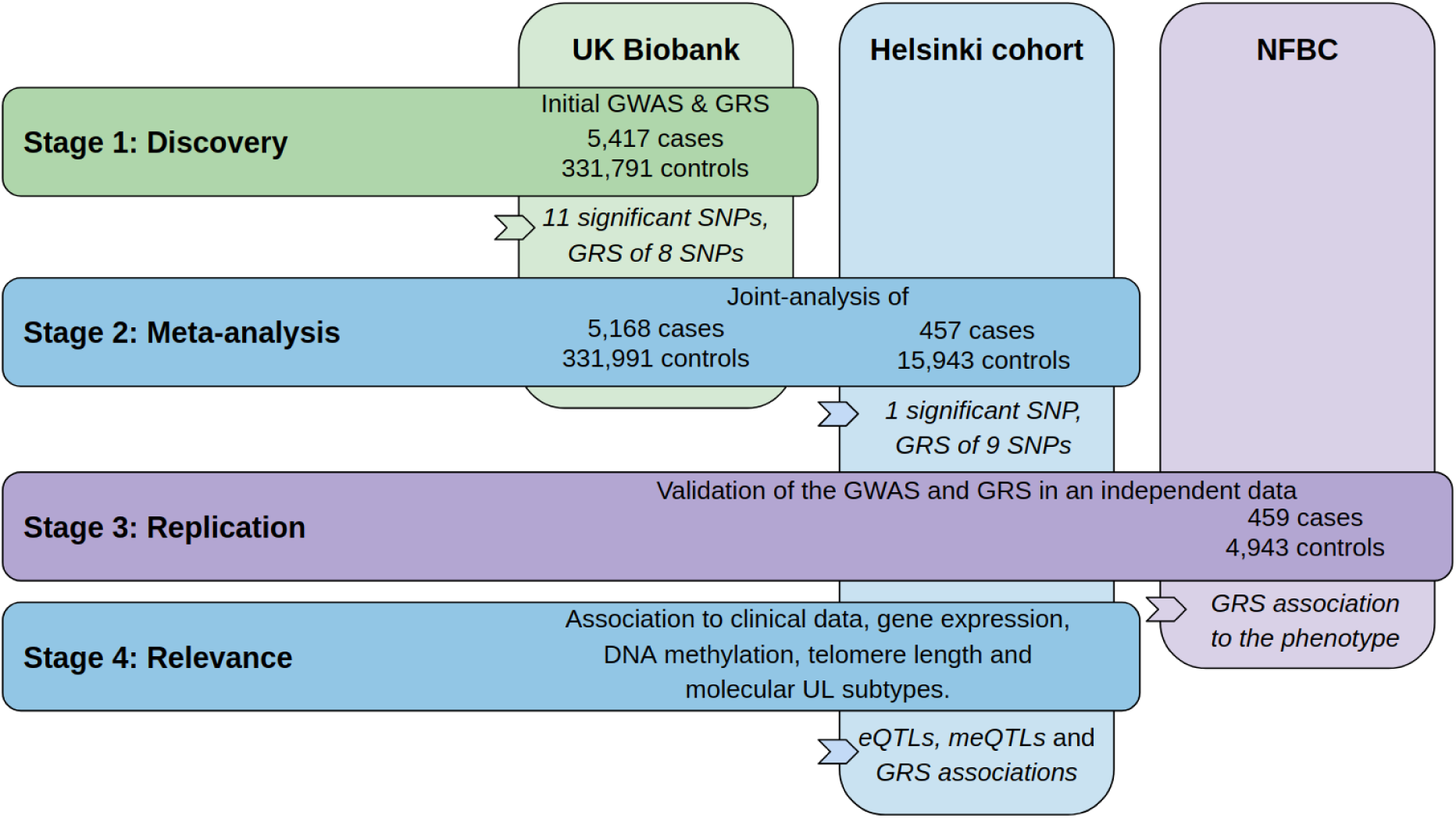
Outline of the study stages and genotyping cohorts. GRS, genomic risk score. NFBC, Northern Finland Birth Cohort.

### Patient and tumor material

Our in-house patient and tumor data were investigated regarding the identified risk loci. Clinical background data for the number of ULs (Fig. S1) and age at hysterectomy (Fig. S2) were available for 357 patients in the Helsinki cohort. All tumors of ≥1cm diameter had been harvested and stored fresh-frozen, and *MED12*-mutation status was screened from all 1,165 tumors. Gene expression was studied in an 1Mb flank from each SNP of interest using 60 tumors and 56 respective normal myometrium samples. DNA methylation was studied in an 1Mb flank using 56 tumors and 36 normals. Telomere length estimates derived from whole genome sequencing data were available for 37 UL and 28 myometrium samples. The study was approved by the ethics review board of the Helsinki University Central Hospital, Finland. Details regarding materials and methods are given in the Supplementary Methods.

### Statistical analysis

For GWAS, P<5×10^−8^ was reported as significant. The GRS association tests (Table S6) were controlled for family-wise error rate (FWER) and reported significant for Holm-Bonferroni adjusted P<0.05. Other families of association tests were controlled for false discovery rate (FDR; Benjamini-Hochberg method) and noted significant at FDR<10%.

## RESULTS

### Identification of predisposition loci

At discovery stage 11 SNPs emerging from five distinct genetic loci passed the genome-wide significance level of 5×10^−8^: 11q22.3 region in *ATM* (top SNP rs1800057; OR=1.49; P<1×10^−16^), 11p13 near *WT1* (rs74911261; OR=1.21; P=1.7×10^−13^), 6q25.2 near *ESR1* (rs11751190; OR=1.17; P=1.4×10^−11^), 1p36.12 near *WNT4* (rs12038474; OR=1.17; P=1.8×10^−10^) and at 22q13.1 in *TNRC6B* (rs12484776; OR=1.14; P=2.5×10^−8^). The sixth most significant association, not reaching genome-wide significance, was observed at *TERT* locus in 5p15.33 (rs2736100; OR=0.91; P=3.6×10^−7^). The association results are given in Table S1, and the lead SNPs are summarized in Table 1.

**Table 1:** Predisposition loci for uterine leiomyoma.

| Chr | Position | Band | rs-id | A | B | MAF | OR | P | Likely disease gene |
| --- | --- | --- | --- | --- | --- | --- | --- | --- | --- |
| 11 | 108,143,456 | q22.3 | rs1800057 | C | G | 0.028 | 1.49 | **1.00E-16** | *ATM* |
| 11 | 32,364,187 | p13 | rs2057178 | G | A | 0.179 | 1.21 | **1.73E-13** | *WT1* |
| 6 | 152,558,197 | q25.2 | rs11751190 | G | A | 0.226 | 1.17 | **1.43E-11** | *ESR1* |
| 1 | 22,403,357 | p36.12 | rs12038474 | G | A | 0.202 | 1.17 | **1.81E-10** | *WNT4* |
| 22 | 40,652,873 | q13.1 | rs12484776 <sup>a</sup> | A | G | 0.245 | 1.14 | **2.45E-08** | ? |
| 5 | 1,286,516 | p15.33 | rs2736100 <sup>b</sup> | C | A | 0.336 | 0.91 | 3.59E-07 | *TERT* |

Fig. 2 displays a regional structure of each locus and the flanking association values, linkage disequilibrium (LD) and genome annotation. Annotation tracks are included for tissue-specific data on open chromatin, topologically associating domains (TAD) and other regulatory features (details in Supplementary Methods).

**Figure 2:**
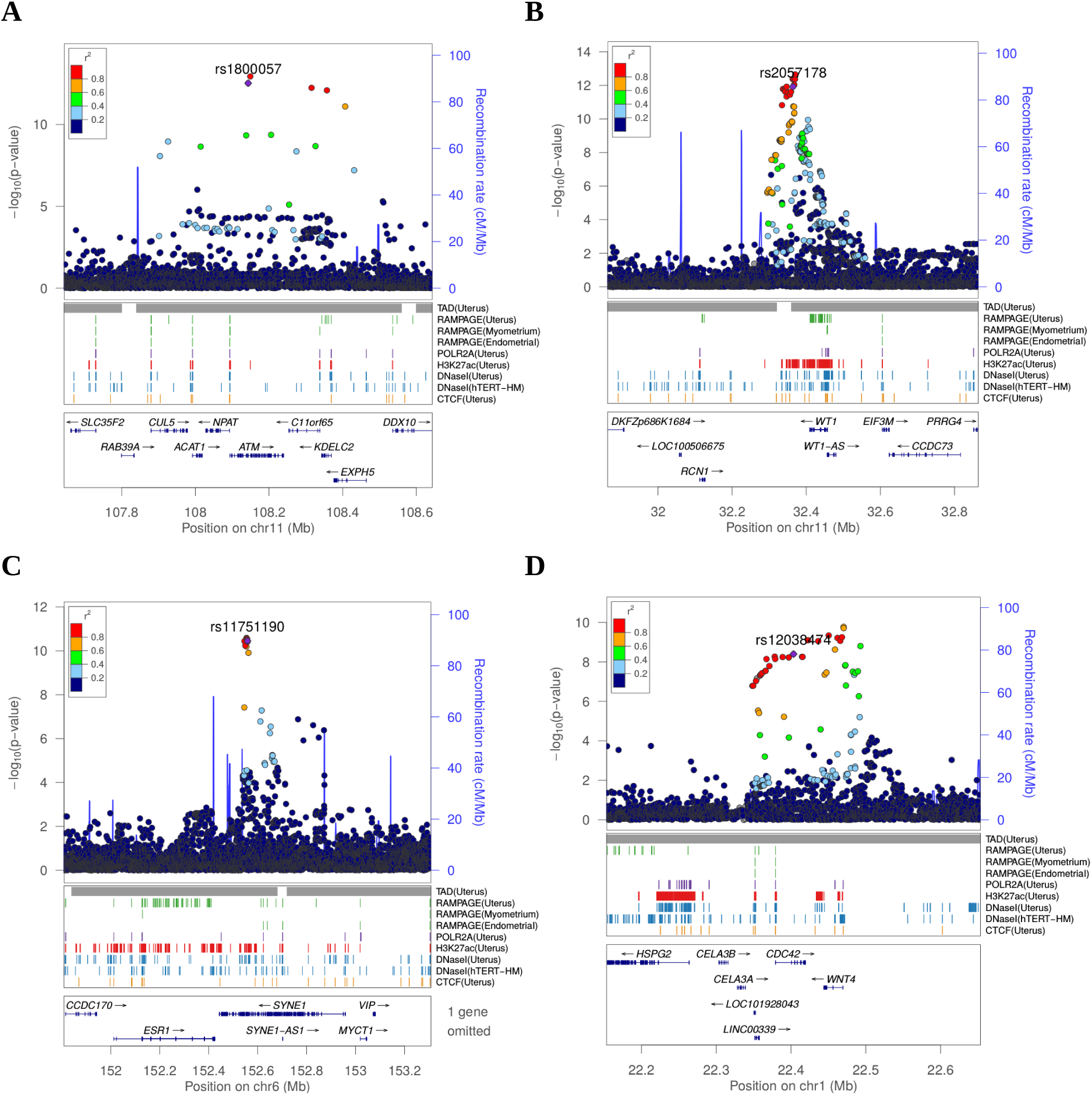
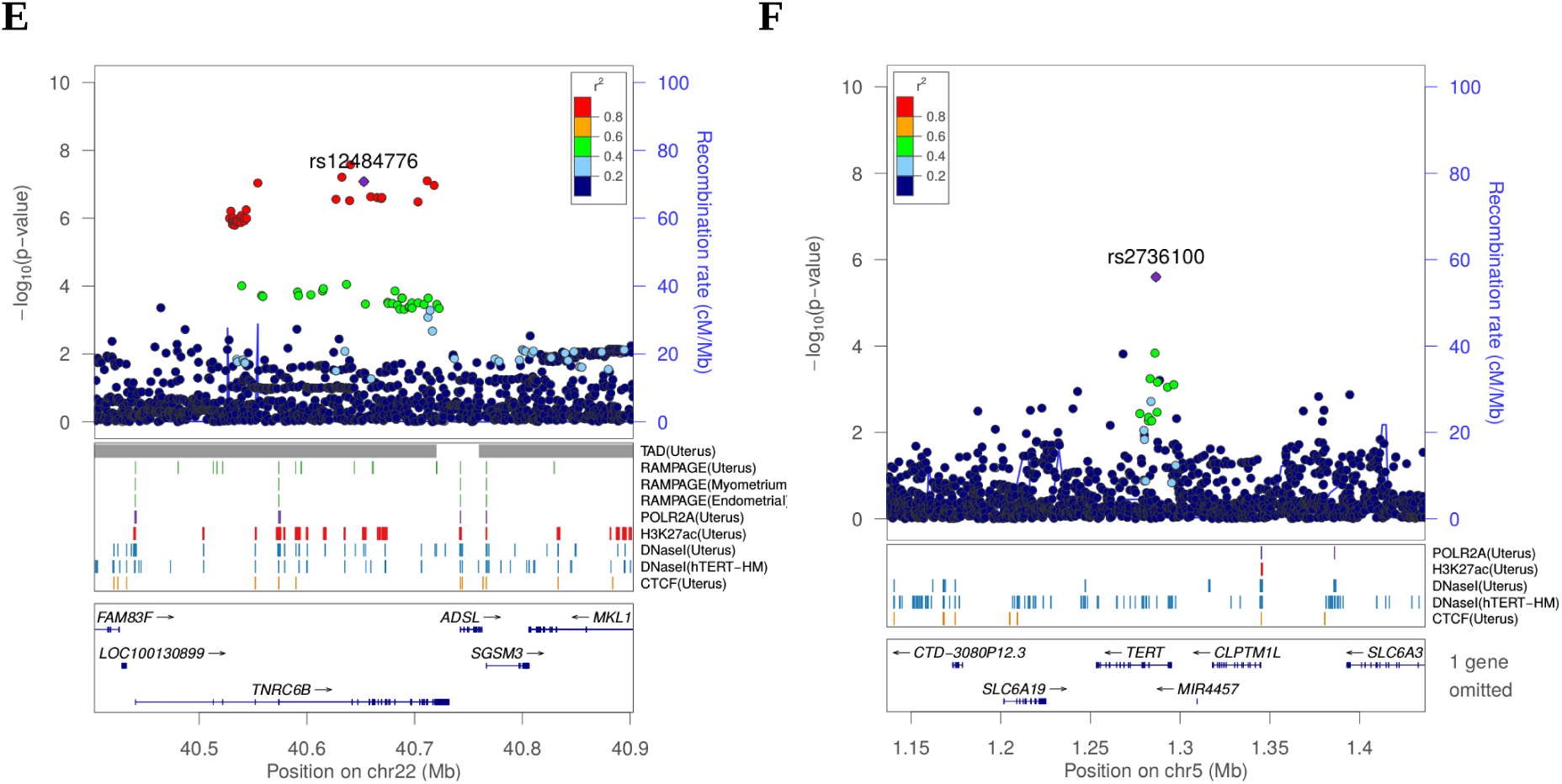
Structure of UL predisposition loci. **A-F**, the SNPs from stage 1 and their genomic context. The LD estimates (r^2^) were taken from UK10k ALSPAC. Associations are based on the genome-wide summary statistics on self-reported UL cases (n=5,168; Neale lab data for 10 million SNPs). Gene symbols and ENCODE tracks (details in Supplementary Methods) are shown for reference; coordinates follow hg19.

### Genomic risk score

A polygenic risk score^20^ was compiled based on the discovery stage associations, including *TERT* locus. After LD thinning (r^2^ <0.3) the 12 discovery-stage SNPs, eight SNPs from the six distinct loci passed for the initial genomic risk score (GRS; Table S3). The SNP weights were based on UKBB log-odds. We applied this initial GRS of eight SNPs to the Helsinki cohort and identified a significant association to the UL phenotype (Wilcoxon rank-sum P=0.0098; adjusted P=0.029; one-sided; W=3.41×10^6^).

### Meta-analysis

The second stage GWAS combined the UKBB and Helsinki cohorts for a meta-analysis approach. We utilized the genome-wide statistics available for a total of 5,168 self-reported UL cases (Neale lab data; details in Supplementary Methods). rs117245733 at 13q14.11 was identified as the only SNP with a suggestive (P<10^−6^) association in both the UKBB (OR=1.34; P=3.31×10^−6^) and Helsinki (OR=1.82; P=8.12×10^−6^) cohorts. The combined association was genome-wide significant (fixed effect model OR=1.41; P=3.20×10^−8^). Fig. 3 shows the regional structure and combined association at the locus: the SNP resides on a gene poor region, at a conserved element that has activity in uterus-specific H3K27ac and DNaseI data (see ENCODE track details in Supplementary Methods). Summary of the meta-analysis results is given in Table S2. The SNP rs117245733 at 13q14.11 - weighted by its UKBB log-odds - was appended to the initial GRS model. The final GRS model of nine SNPs and their UKBB-based weights is given in Table S3.

**Figure 3:**
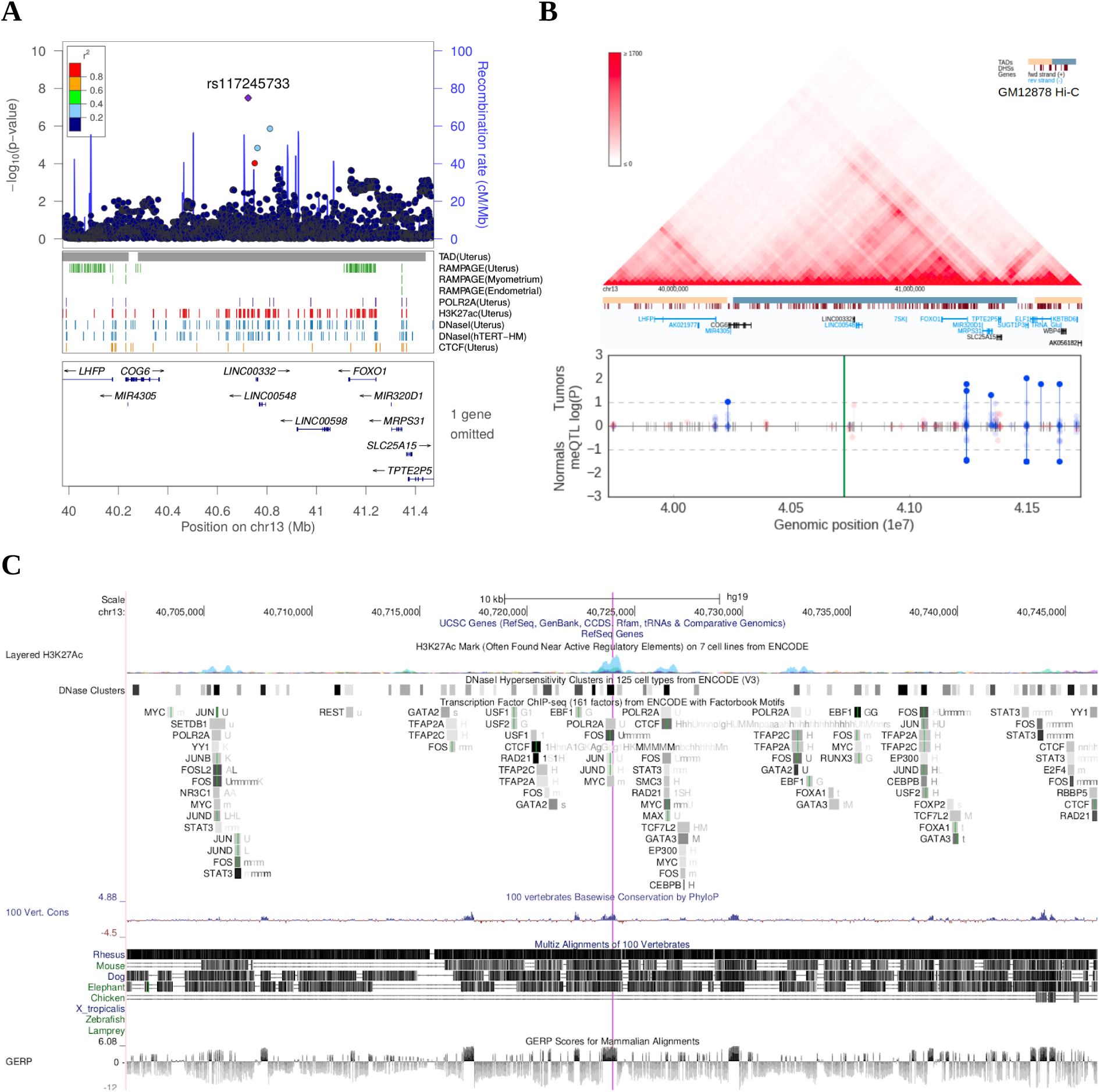
Meta-analysis of UL risk revealed rs117245733 at a gene poor locus at 13q14.11. **A**, meta-analysis P-values and the genomic context at the locus. Gene symbols and ENCODE tracks (details in Supplementary Methods) are shown for reference; coordinates follow hg19. **B**, Hi-C, TADs and CpG methylation around the locus with an 1Mb flank. The needle plot shows the meQTL associations (dashed lines at 10% FDR; green line denotes the SNP; gray ticks denote all CpGs tested; blue needle for positive coefficient, red otherwise) for tumors (above x-axis; n^AA^ =53, n^AB^ =3) and normals (below x-axis; n^AA^ =33, n^AB^ =2). **C**, UCSC genome browser tracks related to conservation and regulation at the locus.

### Replication of the GWAS and GRS

The third stage replicated the observations in NFBC. The SNP identified in the stage 2 meta-analysis, rs117245733 at 13q14.11, was replicated with a nominal P=0.034 (linear mixed model; OR=1.50; 95% CI 1.03-2.19). All SNPs except rs11031736 at 11p13 had the same effect direction as observed in UKBB, however, none of the single-SNP associations in NFBC held with multiple testing (Table S4). The association between the GRS and UL phenotype replicated with a significant P=1.7×10^−5^ (Wilcoxon rank-sum; adjusted P=0.0001; one-sided; W=1.0×10^6^). Analysis of the case-control distributions showed an odds ratio of 2.09 for one-unit increase in GRS values (logistic regression; 95% CI 1.43-3.04; Fig. S4-S5). The GRS model passed the goodness-of-fit test (Hosmer-Lemeshow P=0.406), and the sensitivity and specificity characteristics of the model are summarized as a receiver operating characteristic (ROC; AUC=0.56; 95% CI 0.53-0.59) curve in Fig. S5B.

UL susceptibility is known to vary by ancestry^5^. We computed population specific estimates for the GRS based on the gnomAD database allele frequencies. Fig. S5 shows the resulting estimates for reference (details in Table S7).

### Association to clinical variables

The number of ULs per patient had a significant positive association to the GRS (negative binomial regression P=0.0047; adjusted P=0.024; rate ratio 1.61; 95% CI 1.07-2.44 for one-unit increase in GRS; Fig. S6). No association was found between the GRS and age at hysterectomy (Table S6). Tests for each SNP separately are given in Table S5.

### Association to *MED12* mutated tumors

Our UL set of 1165 lesions included 931 (80%) mutation-positive and 234 mutation-negative tumors. the occurrence of mutant tumors did not distribute evenly among the 357 patients. In total 178 (50%) and 96 (27%) patients had all their tumors identified as either *MED12*-mutation-positive or -negative, respectively (Fig. S7), suggesting that genetic or environmental factors contribute to the preferred UL type in an affected individuals, as previously observed^16^. Indeed, mutation positive patients were found to have a significantly higher GRS (Wilcoxon rank-sum P=0.005; adjusted P=0.024; two-sided; W=6803). This difference in GRS distributions is visualized in Fig. S5.

Strikingly, comparison against the population controls (n=15,943) revealed an opposite effect direction for the above-mentioned patient groups: the MED12-mutation-positive (178) subset of patients displayed an odds ratio of 2.34 for one-unit increase in GRS (95% CI 1.32-4.15) compared to the controls, and the mutation-negative (96) subset an odds ratio of 0.66 (95% CI 0.28-1.54) compared to the controls. Thus, the majority of the compiled case-control association signal had arisen from the *MED12*-mutation-positive subset of the patients.

The number of *MED12*-mutation-positive tumors per patient also had a significant positive association to the GRS (negative binomial model P=0.0026; adjusted P=0.018; rate ratio 2.08; 95% CI 1.13-3.83 for one-unit increase in GRS; Fig. S8). No association between the number of *MED12*-mutation-negative tumors and GRS was found (Fig. S8). Tests for each SNP separately are given in Table S5.

### Association to gene expression

In tumors, one cis expression quantitative trait loci (cis-eQTLs) passed a 10% FDR. The risk allele of rs61778046 at 1p36.12 showed positive correlation with both *WNT4* expression (nominal P=0.004, Fig. 4B) and *alkaline phosphatase, liver/bone/kidney* (*ALPL)* expression (P=3.07×10^−5^, Fig. S11). *WNT4* and *ALPL* are in adjacent TADs (Fig. 4C). In normal myometria, no cis-eQTL passed the 10% FDR. Altogether 11 genes in tumors and 8 genes in normals passed with a nominal P<0.05 (Table S8). All expression results within 1Mb flank can be found in Supplementary Data 1-2.

**Figure 4:**
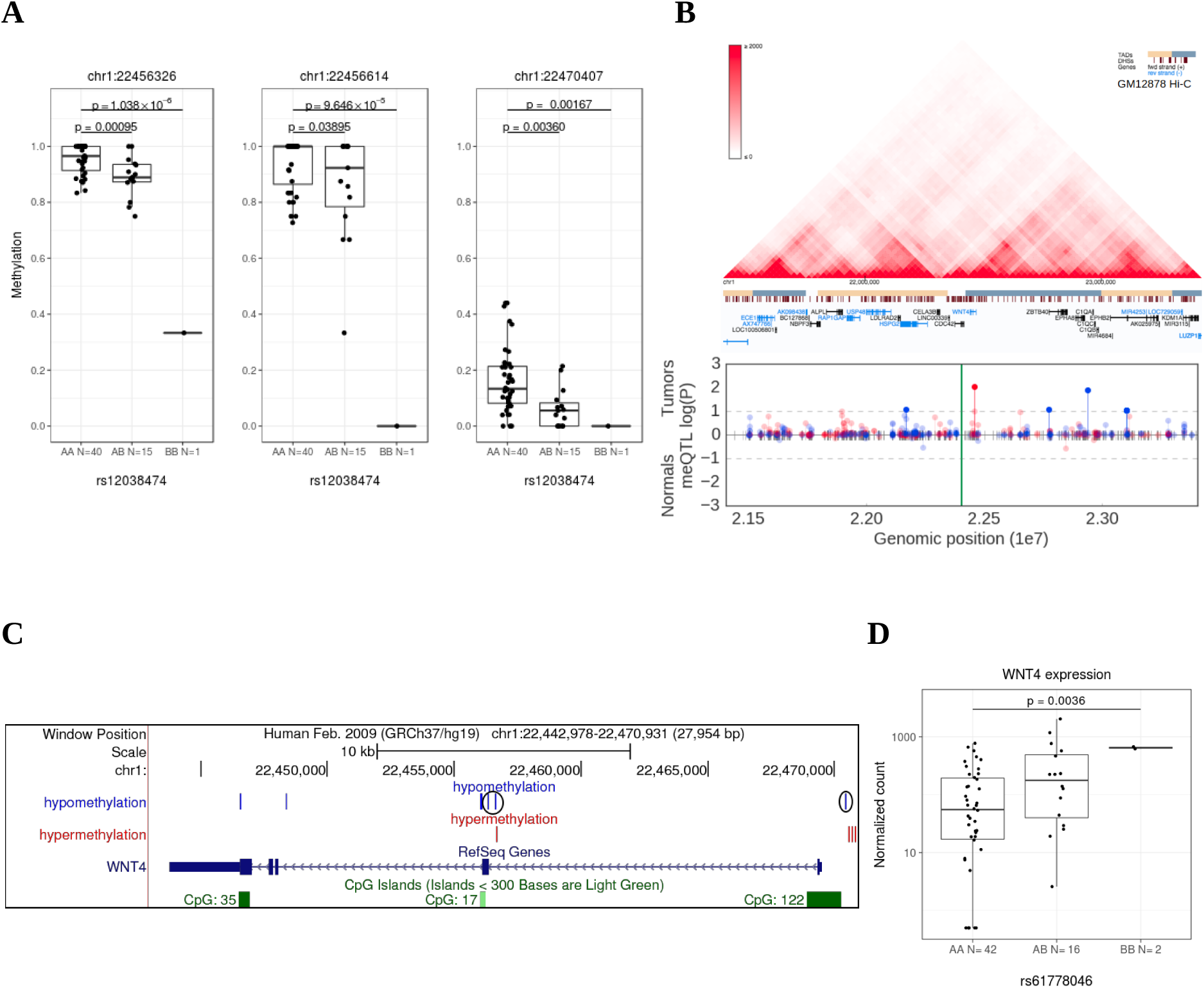
Methylation and expression differences in *WNT4*. **A**, Tumors (n=56) show methylation differences in three meQTLs (chr1:22456326, chr1:22456614 and chr1:22470407). **B**, Hi-C, TADs and CpG methylation around the locus with an 1Mb flank. The needle plot shows the meQTL associations (dashed lines at 10% FDR; green line denotes the two SNPs; gray ticks denote all CpGs tested; blue needle for positive coefficient, red otherwise) for tumors (above x-axis; n^AA^ =40, n^AB^ =15, n^BB^ =1) and normals (below x-axis; n^AA^ =23, n^AB^ =12). **C**, *WNT4* locus. Hypomethylation refers to decreased methylation in BB vs. AA genotype and hypermethylation the opposite. The three CpGs from panel A are shown circled. **D**, Expression differences in tumors (n=41) *WNT4* based on rs61778046 genotype.

Altogether 95 cis-splicing quantitative trait loci (cis-sQTL) had a suggestive correlation (P < 0.05) within 1Mb flank of each SNP. No cis-sQTL passed the 10% FDR. All the cis-sQTLs are listed in Supplementary Data 4.

### Association to DNA methylation

Altogether 3,340 (1,865 in tumors and 1,475 in normals) cis methylation quantitative trait loci (cis-meQTL) had suggestive associations with nominal P < 0.05. Of these, 38 passed a 10% FDR. Of the plausible culpit genes, *TERT*, *WT1* and *WNT4* showed most significant meQTL associations. 18 of the meQTLs in these genes were detectable both in tumors and normals (Table S9). All the cis-meQTLs and annotation for their genomic context are in Supplementary Data 3. The association of selected meQTLs with expression is shown in Fig. S12-S13.

### Association to telomere length

When examining telomere lengths in tumors from carriers and noncarriers of the *TERT* risk allele, the risk allele (rs2736100) was significantly associated with a shorter telomere length (Kruskal-Wallis P=0.01) (Fig. S14). The association was not seen in myometrium. Overall the telomere length was significantly shorter in tumors than normals (P=0.01), as previously reported^21,22^. Adjusting for the patient age did not explain away the association in tumor data. No association was detected between the genotype and the number of somatic structural variants.

### Previously proposed UL predisposition loci

Previous UL association studies^9,13–15^ have reported altogether seven genome-wide significant UL susceptibility loci. Two out of the seven loci - that is, 22q13.1 (at *TNRC6B*) and 11p15.5 (at *BET1L*) - replicated in the UKBB summary statistics for 5,168 self-reported UL cases (Neale lab data). See Table S10 for a summary of these results.

## DISCUSSION

The UK Biobank genotype-phenotype data revealed five novel predisposition loci for UL in close proximity of highly plausible culprit genes. A meta-analysis of the UKBB and Helsinki cohorts identified a sixth novel locus at 13q14.11, residing within a conserved element that has enhancer activity in uterus-specific experiments (Fig. 3). Two previously reported loci, at *TNRC6B^9–12^* and *BET1L^9,23^*, were also validated, and are indeed linked to UL predisposition. For the three latter loci mechanistic connection to UL is obscure at present.

Though simple association is not sufficient to formally prove causality, the five new risk genes implicated in the discovery phase of this study, *ESR1, WT1, WNT4, ATM and TERT,* form a robust set of likely culprits.

Estrogen is a well known inducer of UL growth^24^. The top association at 6q25.2 (rs11751190) resides within intron 23 of *Spectrin Repeat Containing Nuclear Envelope Protein 1* (*SYNE1*), 130kb downstream of *ESR1*, the latter being the only gene that resides completely within the TAD (Fig. 2C). While the role of estrogen in leiomyomagenesis has been firmly established, this is the first genetic evidence to this end.

*WT1* and *WNT4* are central factors in uterine development^25,26^, and perturbations in their function are known to have neoplastic potential. The strongest association at 11p13 (rs2057178) maps to an intergenic region, 45kb downstream of the closest gene *WT1* and the region has active enhancer activity in uterus (Fig. 2B). *WT1* is a transcription factor that acts as both a tumor suppressor and an oncogene^27^. The chromosome 1p36.12 association (rs12038474 and rs61778046) arises at the promoter region of *Cell Division Cycle 42* (*CDC42*) and roughly 50kb downstream of *WNT4*. While *CDC42* cannot be dismissed as a candidate gene, tumor derived data suggests *WNT4* as the more plausible target. The risk allele at rs61778046 was associated with upregulation of *WNT4* (Fig. 4B). The effect is also supported by GTEx Portal data, where *WNT4* shows suggestive upregulation associated with the presence of risk allele in myometrium (P=0.044; accessed Dec 1, 2017). *WNT4* is known to be overexpressed in uterine leiomyomas with *MED12* mutations^28^. Taken together the risk allele may cause increased expression already in the normal myometrium, and the effect could be further selected for during UL genesis. The risk locus in 1p36.12 was also associated with several meQTLs suggesting that methylation may have a role in *WNT4* regulation (Fig. 4). *WNT4* encodes a signaling protein that has a crucial role in sex-determination^29^, and the WNT signaling pathway has a well-established role in various malignancies such as breast and ovarian cancer^30^. Of note, recent GWAS on gestational duration suggested that binding of the estrogen receptor at *WNT4* is altered by rs3820282 (r^2^ =0.9 with our lead SNP)^31^.

*ATM* and *TERT* could be involved in uterine neoplasia predisposition through genetic instability. The lead SNP at 11q22.3 is a nonsynonymous variant (rs1800057; p.Pro1054Arg) in exon 22 of *ATM*. This SNP was recently highlighted in a renal cell carcinoma GWAS^32^. *ATM* is involved in DNA damage response^33^ and is one of the relatively few genes that have been found to be recurrently mutated in leiomyosarcoma^34^. *TERT* encodes a subunit of the telomerase enzyme, which guards chromosomal stability by elongating telomeres. It is expressed in germ cells as well as in many types of cancers^35^. The risk allele at the *TERT* intron 2 (5p15.33; rs2736100) has several associated cis-meQTLs in the 1Mb flank region (Fig. S9-S10). For example CpG chr5:1277576 in the sixth intron of *TERT* and CpG chr5:1285974 in the second intron of *TERT* are detectable both in tumors and normals with nominal P<0.001. While the neoplasia predisposing effect of this SNP is overwhelmingly documented, previous studies have reported contradicting observations on its effect on telomere length^36–40^. ULs have been shown to display shortened telomeres^21,22^, potentially provoking chromosomal instability as the chromosome tips are worn out. Our data shows that this finding is explained by *TERT* risk allele carrier status (Fig. S14).

GRS associated merely with a susceptibility to the most common UL subtype, *MED12* mutation positive tumors. Indeed it has been known that *MED12*-mutation-positive tumors do not distribute randomly among patients^16^, and our data provide at least a partial explanation to this intriguing finding. It may be that environmental factors contribute more significantly to genesis of *MED12* wild type lesions. In our recent study this tumor type was associated with history of pelvic inflammatory disease, and thus infectious agents could be one underlying factor^41^. Obviously, also the power of GWAS to detect genetic associations to more rare UL types is reduced. Much additional work is needed to fully elucidate the mechanisms connecting the risk alleles and emergence of *MED12* mutant UL.

This work highlights several new genetic cornerstones of UL formation, and represents another step towards a much improved understanding of its molecular basis. The GRS score can stratify the female population to low-risk and high-risk quartiles that differ two-fold by their UL risk. While the increased risk appears minor on individual level, the population-level burden to women’s health arising from these risk loci is highly significant considering the incidence of the condition. Together with the recent progress in molecular tumor characterization and subclassification, the identification of the genetic components of UL predisposition should pave the way towards more sophisticated prevention and management strategies for these extremely common tumors. The risk SNP with the most immediate potential value is that at estrogen receptor alpha, and our findings should fuel much further work on the interplay between individual germline genetics, endogenous and exogenous hormonal exposure, and occurrence and growth rate of UL.

## ACKNOWLEDGEMENTS

We are thankful to Sini Marttinen, Sirpa Soisalo, Marjo Rajalaakso, Inga-Lill Åberg, Iina Vuoristo, Alison Ollikainen, Elina Pörsti, Salla Välipakka and Heikki Metsola for their technical support. We also thank Pirjo Ikonen and the rest of the staff of the Kätilöopisto Maternity Hospital, and the staff of the Department of Pathology, University of Helsinki for technical assistance. We thank Minna Männikkö, Tuula Ylitalo, and the rest of the staff of the Northern Finland Birth Cohort Studies, Faculty of Medicine, University of Oulu for technical assistance. The study was supported by grants from Academy of Finland (Finnish Center of Excellence Program 2012-2017, No. 1250345), European Research Council (ERC, 695727), Cancer Society of Finland, Sigrid Juselius Foundation and Jane and Aatos Erkko Foundation. NV received a grant from the Academy of Finland (No. 287665). KP received a grant from Nordic Information for Action eScience Center (NIASC), the Nordic Center of Excellence financed by NordForsk (No. 62721). The authors would like to thank the Rivas lab for making the GBE resource available. We thank the Neale lab for making the UKBB summary statistics available.

## AUTHOR CONTRIBUTIONS

AP, OH, JS, RB and NS collected the tissue samples. NV and HK analyzed the data and wrote the first draft of the manuscript. HRH, JT, SB, TT, JA, TP, AA, RL and EK participated in the data collection and analysis. LAA and KP supervised the study. All authors reviewed the final draft.

